# Electrostatic Trade-Off between Mesophilic Stability and Adaptation in Halophilic Proteins

**DOI:** 10.1101/2024.01.08.574673

**Authors:** Pablo Herrero, Alba Pejenaute, Oscar Millet, Gabriel Ortega

## Abstract

Extremophile organisms have adapted to extreme physicochemical conditions. Halophilic organisms, in particular, survive at very high salt concentrations. To achieve this, they have engineered the surface of their proteins to increase the number of short, polar and acidic amino acids, while decreasing large, hydrophobic and basic residues. While these adaptations initially decrease the thermodynamic stability in the absence of salt, they grant halophilic proteins remarkable stability in environments with extremely high salt concentrations, where non-adapted proteins unfold and aggregate. The molecular mechanisms by which halophilic proteins achieve this, however, are not yet clear. Here, we test the hypothesis that the halophilic amino acid composition destabilizes the surface of the protein, but in exchange improves the stability in the presence of salts. To do that, we have measured the folding thermodynamics of various protein variants with different degrees of halophilicity in the absence and presence of different salts, and at different pH values to tune the ionization state of the acidic amino acids. Our results show that, although electrostatic interactions decrease the stability of halophilic proteins, in exchange they induce a significant salt-induced stabilization and improve solubility. Besides electrostatic interactions, we also show that other general contributions, such as hydrophobic effect and preferential exclusion, are important. Overall, our findings suggest a trade-off between folding thermodynamics and halophilic adaptation to optimize the stability of halophilic proteins in hypersaline environments.

**Significance statement:** This work explores how extreme halophiles adapt their proteins for survival in hypersaline environments. By engineering the protein surface, evolution has selected proteins adapted to high salt concentrations. Our findings suggest a delicate balance between protein stability and haloadaptation modulated in part by electrostatic interactions, furthering our understanding of life adaptation to extreme environments.

## Introduction

Life has adapted to many different habitats on Earth, including environments with extreme conditions of pressure, temperature, and chemical composition. As opposed to mesophiles, extremophiles have developed multiple adaptations to survive to the extreme physicochemical challenges imposed by these environments. Piezophiles living at the bottom of deep-sea trenches, for example, have evolved lipid compositions that preserve the integrity of their cellular membranes at pressures orders of magnitude higher than those at sea level (Winnikoff et al., 2021). Likewise, hyperthermophiles adapted to hydrothermal vents and hot springs have evolved proteins with increased thermostability to cope with extremely high temperatures (Robic et al., 2003).

Halophiles are a class of extremophilic organisms that have adapted to environments with very high salinity (Oren, 2006). The main challenge at these conditions arises due to the high osmotic stress imposed by the extracellular salt concentration, which drains the water out of the cytoplasm and desiccates the cell. To prevent this, halophilic organisms have evolved two different mechanisms depending on the degree of adaptation required. Halotolerant organisms, for example, can withstand moderate extracellular salt concentrations up to 1 M. To compensate the external osmotic pressure and maintain the water content within the cytoplasm, they synthetize and accumulate large intracellular amounts (up to hundreds of millimolar) of organic cosolutes (Yancey et al., 1982). These compounds, traditionally termed compatible solutes, preserve, and sometimes even improve, the solubility and stability of proteins (Pais et al., 2009; Graziano, 2012). Examples of these cosolutes include polyols such as sucrose and trehalose, but also amino acids such as glycine, betaine, sarcosine and taurine (Yancey, 2005). The synthesis of these compounds, however, demands a large amount of mass and energy, and is thus not suitable for higher salt concentrations (Ding et al., 2022).

Adaptation to more extreme salt concentrations requires a different strategy. Extreme halophiles, such as halophilic archaea (Oren, 2006) and some halobacteria (Antón et al., 2002), thrive in environments where extracellular salt concentrations exceed 1 M, sometimes even approaching saturation levels. To achieve this, extreme halophilic archaea drastically increase salt concentration in their cytoplasm. By upregulating potassium-selective ion channels (Meury and Kohiyama, 1989; Kraegeloh et al., 2005), they accumulate intracellular concentrations of potassium chloride even above 3 M (Christian and Waltho, 1962). This extremely high cytoplasmic salt concentration, however, poses a significant challenge to the normal functioning of the cellular machinery.

Extreme halophilic organisms have adapted their proteome to remain functional at the extremely high salt concentrations of their cytoplasm. High molar salt concentrations usually result in irreversible unfolding and aggregation of mesophilic proteins (Dennis and Shimmin, 1997). To prevent this, proteins from extreme halophiles feature a unique amino acid composition, disfavoring large, bulky and hydrophobic residues, while favoring short, polar and negatively charged residues (Madern et al., 2000; Paul et al., 2008). Most of this reengineering occurs at the surface, with lysine-to-glutamate and lysine-to-aspartate amongst the most common substitutions (Fukuchi et al., 2003). There are also, however, modifications inside the hydrophobic core of the protein, such as leucine-to-valine and isoleucine-to-valine substitutions (Paul et al., 2008). As a result of this characteristic signature in their amino acid composition, halophilic proteins manage to remain soluble and functional at high molar salt concentrations (Ebel et al., 1999; Polosina et al., 2002), while displaying structures and functions highly conserved with respect to their non-adapted homologues (Pica et al., 2013; Graziano and Merlino, 2014).

Efforts to understand haloadaption have mostly focused on surface electrostatics. Motivated by the unusual abundance of acidic amino acids on the surface of halophilic proteins, traditional models have explained halophilic adaptation almost exclusively by the stabilizing effect of ionic interactions between clusters of carboxylic acids on the surface of halophilic proteins and potassium cations in solution, either through direct carboxylate-potassium interactions or through the formation of a tight water-potassium hydration shell (Zaccai et al., 1989; Mevarech et al., 2000; Ebel et al., 2002). More recent research, however, has shown that the stabilization of halophilic proteins correlates better with the composition of the solvent-exposed hydrophobic area than with the charge content of the protein (Tadeo et al., 2009; Tadeo et al., 2009; Ortega et al., 2015). This experimental evidence points to an alternative mechanism, by which the halophilic signature in amino acid composition reduces apolar content at the protein surface, thus modulating the hydrophobic effect to improve protein stability at high salt concentrations.

In this work, we set out to explore quantitatively the effect of electrostatic interactions in the stability of halophilic proteins. To do that, we have measured the stability of halophilic and non-halophilic model proteins, modulating the pH to control the ionization state of carboxylic groups, and in the absence and presence of different salts. Our results allow us to quantify the effect of electrostatics on the stability of halophilic proteins, and the extent to which these interactions are important in halophilic adaptation. More broadly, our results also help us better understand protein-cosolute interactions, and the different mechanistic contributions involved.

## Results

We have studied a set of three protein variants with different degrees of halophilicity, with non-halophilic, intermediate, and extreme halophilic amino acid compositions. As our non-halophilic model, we have studied protein L, a 64-amino acid protein from the mesophilic bacteria *Finegoldia magna* with well-characterized thermodynamics (Wikström et al., 1994). In addition, we have also studied two protein L variants with different degrees of halophilicity, which we had previously developed by systematically engineering their amino acid composition to gradually resemble that typical of halophilic proteins (Tadeo et al., 2009). More specifically, given that lysine-to-glutamate substitutions are one of the most abundant in halophilic proteins, we have studied a protein L variant in which five of its seven surface-exposed lysines L are substituted for glutamates. This variant features a much higher negative charge (its net charge is -13, compared to -3 of protein L) and less solvent-exposed area than the non-halophilic wild type protein L, thus being a suitable model for an extreme halophilic protein. Finally, we have also studied a variant with intermediate halophilic character, in which we substituted the same five lysines for glutamines instead. This variant features a net negative charge of -8, intermediate between the extreme halophilic and the non-halophilic variants, and solvent-exposed area composition similar to the extreme halophilic variant, thus being a good model for a halotolerant protein. The high structure and sequence homology of these three protein variants will help us identify effects arising due to halophilic amino acid composition, separating these from more protein-specific effects.

We have used these three model proteins to explore systematically the mechanistic contributions to halophilic adaptation. By measuring protein stabilities in the absence and in the presence of increasing concentrations of salt, we have evaluated the mesophilic stability (that is, the stability at low salt concentrations) as well as the effect of different salts on the stability for our three model proteins. More specifically, we have performed chemical denaturation experiments with guanidinium chloride, which we have monitored by circular dichroism, to calculate protein unfolding free energies (Santoro and Bolen, 1988). While we had previously evaluated the effect of salt on the thermostability of halophilic proteins by means of thermal denaturations, the more rigorous characterization of the folding thermodynamics we perform here allows us to identify more accurately the different mechanistic contributions involved. For all three proteins and at all the conditions tested in this work, we find that our chemical denaturation data shows, as expected for protein L, a cooperative unfolding transition with no intermediates (Scalley et al., 1997) (Fig. 1). We can therefore fit our data to the equations for a two-state folding equilibrium to determine protein unfolding free energies (Eqs. 1 and 2; see experimental methods section for more details).

**Figure 1.**
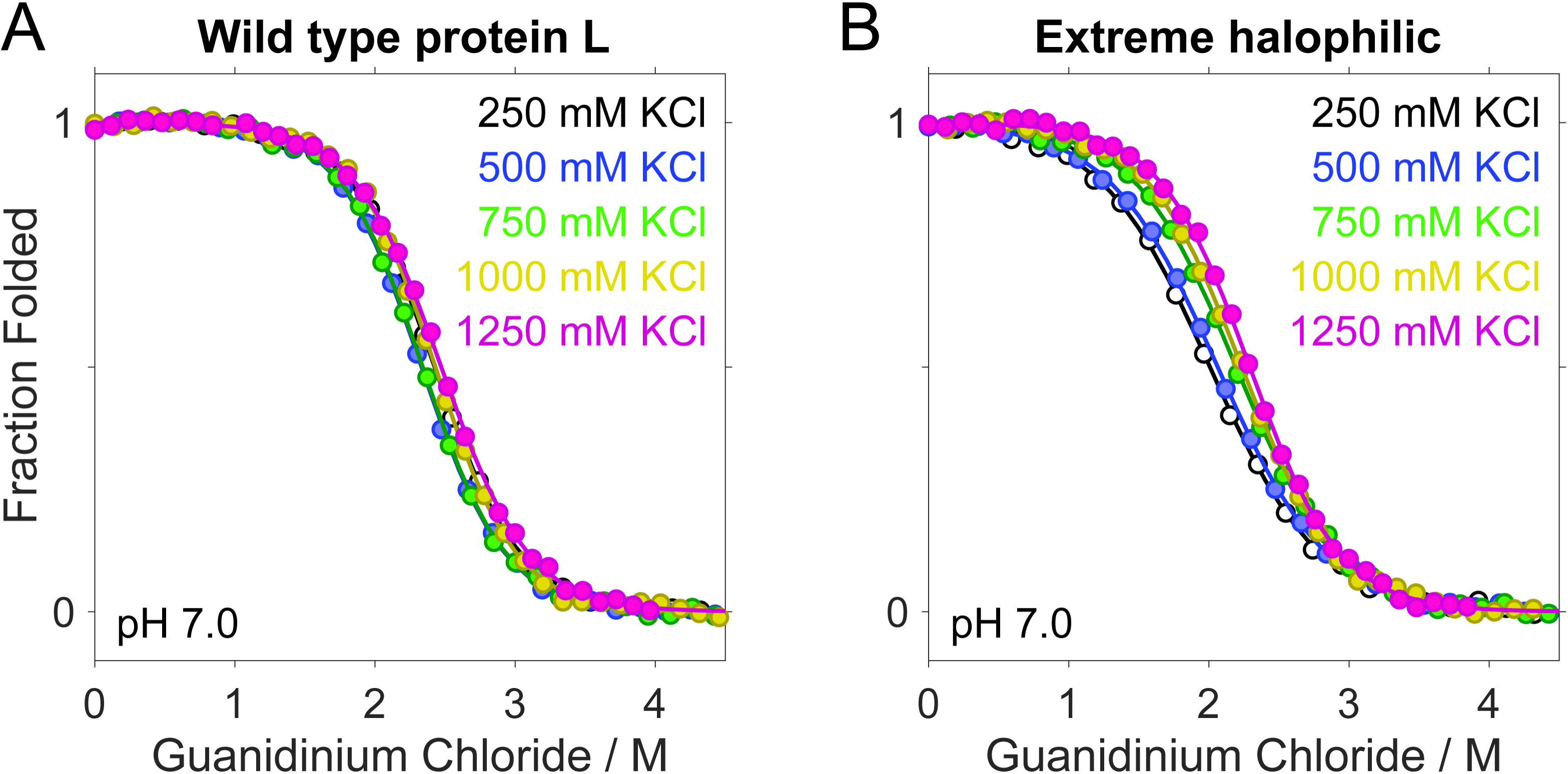
Chemical denaturation experiments to measure the effect of salts on protein stability. (**A**) Overlay of guanidinium chloride-induced denaturation experiments of non-halophilic wild type protein L at increasing concentrations of potassium chloride shows no effect on its folding thermodynamics. (**B**) In contrast, addition of potassium chloride increases the stability of the extreme halophilic lysine-to-glutamate variant, as shown by the shift of the midpoint of the denaturation towards higher guanidinium chloride concentrations.

To tease apart the contribution of electrostatic interactions, we have performed our experimental characterization at two different pH values. First, we have measured protein stabilities at pH 7.0. At this neutral pH, all the carboxylic groups are fully deprotonated and negatively charged, thus being able to form electrostatic interactions with the cations in solution. Then, we have performed the same measurements at a pH low enough to ensure full protonation of all the acidic groups of the protein, thus ablating electrostatic carboxylate-cation interactions. Comparing the results at neutral and low pH will inform us on the role of electrostatic interactions on salt-induced protein stabilization and, therefore, in halophilic adaptation.

We have initially evaluated the mesophilic stability of our three model proteins in the absence of salt (Table 1). In doing this, we find a mesophilic stability of 19.4 ± 0.4 kJ mol^-1^ for the non-halophilic wild type protein L, a value significantly higher than the 10.9 ± 0.5 kJ mol^-1^ measured for the extremely halophilic lysine-to-glutamate variant. In turn, the stability of 15.6 ± 0.5 kJ mol^-1^ measured for the halotolerant lysine-to-glutamine variant is intermediate between the two. We therefore observe a correlation between mesophilic stability and the degree of halophilicity, with more halophilic proteins showing lower stability.

**Table 1.** Comparative analysis of the effect of potassium chloride on the stability of protein L and its halophilic variants at pH 7.0 and 2.6.

| | pH | Mesophilic stability<br>$\Delta G_U^0$ (no salt) / $\text{kJ mol}^{-1}$ | Haloadaptation<br>$m_{KCl}$ / $\text{kJ mol}^{-1} \text{ M}^{-1}$ |
| --- | --- | --- | --- |
| **Wild type protein L** | 7.0 | $19.4 \pm 0.4$ | $0.3 \pm 0.5$ |
| | 2.6 | $16.1 \pm 0.6$ | $0.8 \pm 0.5$ |
| **Extreme halophilic** | 7.0 | $10.9 \pm 0.5$ | $5.3 \pm 0.6$ |
| | 2.6 | $12.5 \pm 0.6$ | $3.4 \pm 0.6$ |
| **Halotolerant** | 7.0 | $15.6 \pm 0.5$ | $1.9 \pm 0.5$ |
| | 2.6 | $15.1 \pm 0.6$ | $2.0 \pm 0.5$ |

Uncertainties reported here and elsewhere correspond to 95% confidence intervals.

To determine the extent to which electrostatics influence the mesophilic stability of halophilic proteins, we have explored protein stabilities at low pH (Table 1). To do this, we have measured the stability of the three proteins in a glycine-hydrochloric acid buffer at a pH of 2.6. This pH value is low enough to ensure that all carboxylic acids are over 90% protonated, as determined by prior NMR studies measuring the values for the microscopic acidity constants for each individual carboxylic acid in protein L (Ortega et al., 2015). Under these conditions, we find that the mesophilic stability of wild type protein L decreases by 3.3 ± 0.7 kJ mol^-1^ with respect to neutral pH. In contrast, the stability at low pH of the extreme halophilic variant increases by 1.6 ± 0.8 kJ mol^-1^, whereas the stability of the intermediate halophilic protein remains unchanged. We attribute these differences to the electrostatic interactions at the surface of the protein. More specifically, for wild type protein L, which has a well-balanced charge content, neutralizing the negative charges of the carboxylic groups increases the electrostatic repulsion between the seven positively charged lysines, thus destabilizing the protein. For the extreme halophilic protein, in contrast, neutralizing the carboxylic groups eliminates the electrostatic repulsion between the (no longer negatively charged) acidic amino acids. Consistent with this, protonation of the carboxylic groups has a negligible effect on the stability of the intermediate halophilic protein.

We have then evaluated the effect of potassium chloride, the most abundant salt in the cytoplasm of extreme halophilic organisms, on haloadaptation (Table 1). At neutral pH, we find that the stability of wild type protein L is only weakly sensitive to the concentration of potassium chloride (Fig. 2A). This is consistent with our expectations for protein L, a non-halophilic protein, as well as with prior reports (Tadeo et al., 2009; Ortega et al., 2015). In contrast, potassium chloride has a much more significant effect on the stability of halophilic proteins. In particular, we find that potassium chloride increases the stability of the extreme halophilic variant, and it does so following the linear dependence on salt concentration expected for salts in the Hofmeister series (Hofmeister, 1888; Baldwin, 1996) (Fig. 2B). We can thus fit our data to a linear equation (Eq. 3) and use its slope (*m_KCl_*) to quantitatively evaluate the stabilizing effect of potassium chloride. At neutral pH, we find a strong stabilizing effect of 5.3 ± 0.6 kJ mol^-1^ M^-1^. This is consistent with prior reports for halophilic proteins (Tadeo et al., 2009; Ortega et al., 2015), and with our hypothesis that this variant is a good model for an extreme halophilic protein. Finally, for our halotolerant variant we find that the effect of potassium chloride on protein stability at neutral pH is, at 1.9 ± 0.5 kJ mol^-1^ M^-1^, intermediate between that seen for the non-halophilic and extreme halophilic proteins (Fig. 2C).

**Figure 2.**
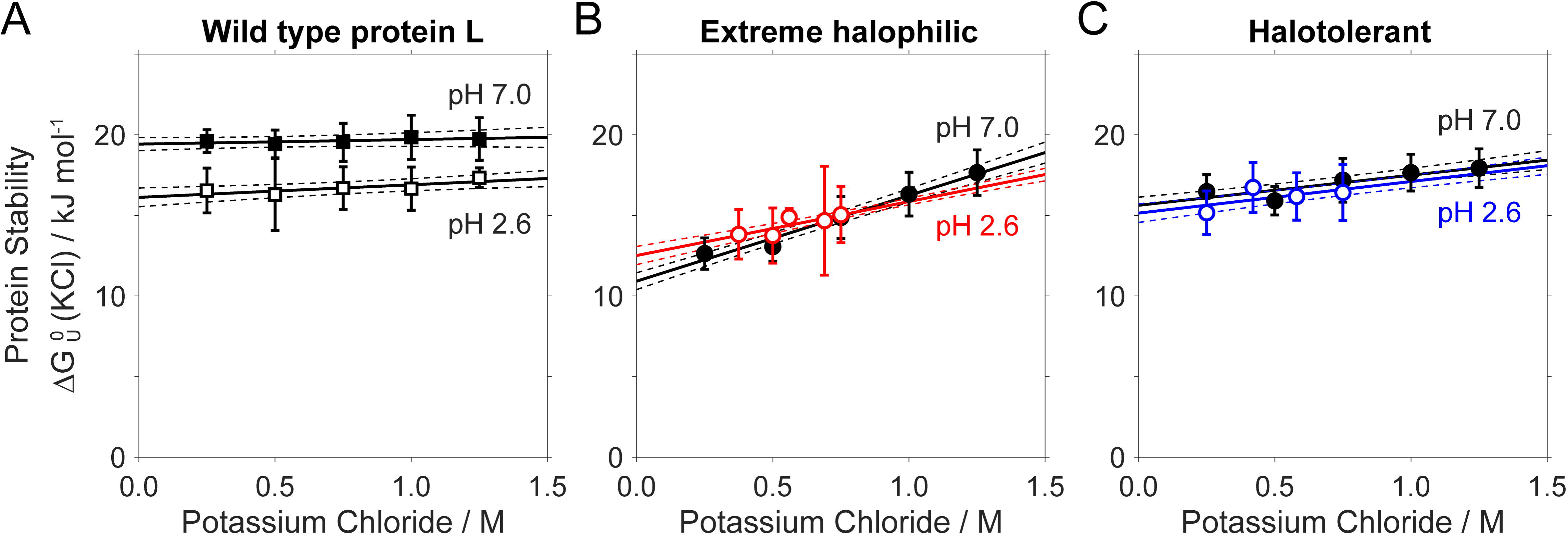
Effect of potassium chloride on the stability of halophilic proteins. (**A**) The mesophilic stability of the non-halophilic wild type protein L at pH 2.6 (open squares) is lower than that at pH 7.0 (black squares), likely because neutralization of the carboxylates increases the repulsion between the positively charged lysines. As expected for a non-halophilic protein, its stability is unaffected by potassium chloride both at neutral and acidic pH (black lines). (**B**) In contrast, the mesophilic stability of the extreme halophilic variant at low pH (open red circles) is higher than that at neutral pH (black circles), due to the decrease in electrostatic repulsion between the carboxylic acids. Consistent with this, potassium chloride increases protein stability at neutral pH (black line) and, to a lesser extent, also at low pH (red line). (**C**) Finally, the mesophilic stability of the intermediate halophilic protein, as well as the effect of potassium chloride, is the same both at neutral (black circles and black line) and acidic (open blue circles and blue line) pH, and intermediate between the ones measured for the non-halophilic and the extreme halophilic proteins. Here and elsewhere in this work, squares and circles represent experimental values, and errorbars 95% confidence intervals derived from fitting the chemical denaturations to a model for a two-state folding equilibrium. Solid lines represent the least-squares linear regression, and dashed lines represent 95% confidence intervals derived from propagating the uncertainties on the experimental stabilities (see materials and methods for more detailed information).

We have next measured the effect of potassium chloride at low pH, to evaluate the extent to which electrostatic interactions are important for the stabilization of halophilic proteins. In doing this we find that, like our results at neutral pH, the effect of potassium chloride in the stability of non-halophilic protein L is, at 0.8 ± 0.5 kJ mol^-1^ M^-1^, again almost negligible (Fig. 2A). This is also consistent with our expectations, given that wild type protein L shows a well-balanced charge content, and thus we do not expect cation-carboxylate interactions to play a significant role. Likewise, the effect of potassium chloride on the halotolerant protein is, at 2.0 ± 0.5 kJ mol^-1^ M^-1^, within error of that measured at neutral pH (Fig. 2C). Protonation of the carboxylic groups thus does not affect the effect of potassium chloride on the stability of the intermediate halophilic protein. However, we observe significant differences when we evaluate the effect of potassium chloride on the extreme halophilic variant. Specifically, we find a stabilizing effect on the extreme halophilic protein that, at 3.4 ± 0.6 kJ mol^-1^ M^-1^, is 1.9 ± 0.8 kJ mol^-1^ M^-1^ lower than that measured at neutral pH (Fig. 2B). This result shows that, while a significant part of the stabilizing effect of potassium chloride in halophilic proteins arises due to electrostatic interactions between carboxylates and potassium cations, this contribution alone is not enough to explain the full extent of the effect of potassium chloride on protein stability, and thus neither halophilic adaptation.

To test whether these observations are general, we also evaluated the effect of sodium sulfate on protein stability. In contrast to potassium chloride, which is a weak stabilizer, sulfate is one of the most stabilizing salts in the Hofmeister series (Ramos and Baldwin, 2002; Tadeo et al., 2009). When we measure the effect of sodium sulfate on the stability of our three model proteins at neutral pH, we observe that the intrinsic stabilities in the absence of salt are within the error of those obtained from the experiments with potassium chloride, adding to the consistency of our experimental characterization (Table 2). As expected for sulfate, the salt-induced stabilizing effects observed are higher than those of potassium chloride for all the three proteins. We also find that its stabilizing effect is more pronounced for the extreme halophilic protein (Fig. 3). Given that the stabilizing effect of sulfate arises mainly due to its tendency to exclude itself from the protein surface (Tadeo et al., 2009), and not due to electrostatic interactions, our results support that halophilic amino acids also optimize other protein properties besides charge composition.

**Figure 3.**
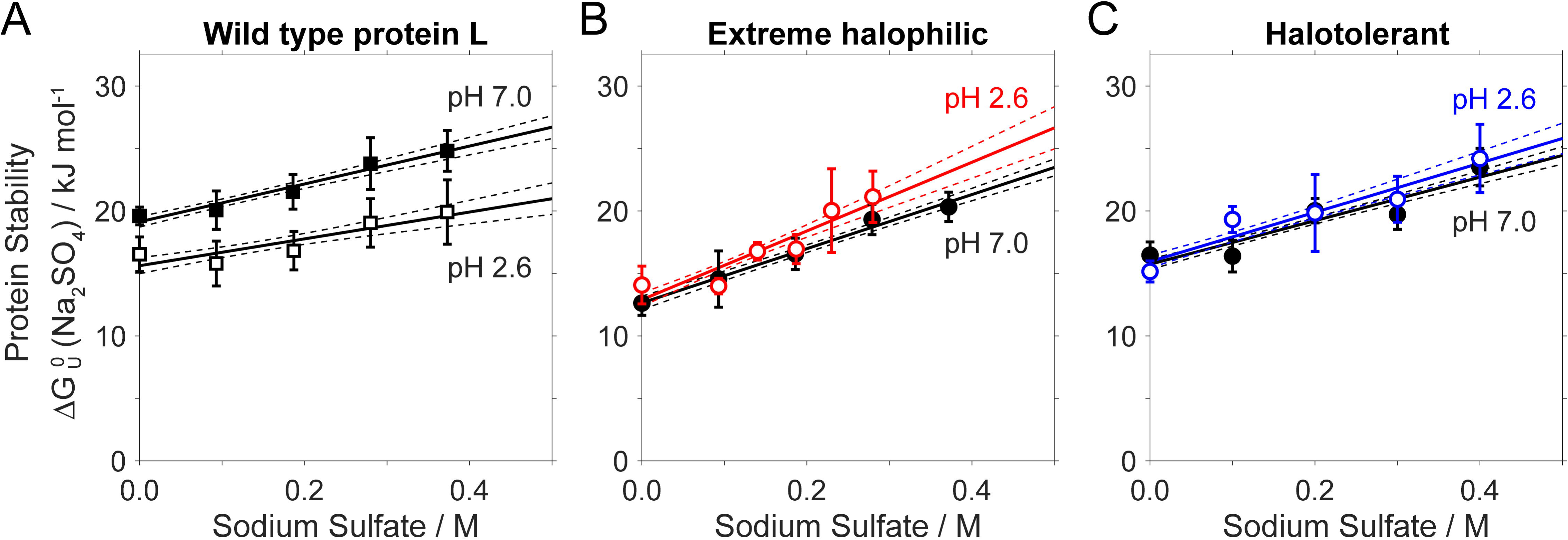
Effect of sodium sulfate on the stability of halophilic proteins. (**A**) As expected for a strong stabilizing salt in the Hofmeister series, sodium sulfate shows a much more important stabilizing effect on non-halophilic protein L than that observed for potassium chloride, and identical at neutral (black squares) and low (open squares) pH. (**B**) The stabilizing effect for the extreme halophilic protein at neutral pH (black circles and line) is also more important than that measured for the non-halophilic protein, but it is even higher at low pH (open red circles and red line), possibly because neutralization of the carboxylates makes the surface of the protein more accessible to the sulfate anions. (**C**) Finally, the stabilizing effect of sulfate on the intermediate halophilic variant is intermediate between those measured for the non-halophilic and extreme halophilic proteins, and identical at neutral (black circles and line) and acidic (open blue circles and blue line) pH.

**Table 2.** Comparative analysis of the effect of sodium sulfate on the stability of protein L and its halophilic variants at pH 7.0 and 2.6.

|  | pH | Mesophilic stability | Haloadaptation |
| --- | --- | --- | --- |
| | | $\Delta G_U^0$ (no salt) / kJ mol<sup>-1</sup> | mNa<sub>2</sub>SO<sub>4</sub> / kJ mol<sup>-1</sup> M<sup>-1</sup> |
| **Wild type protein L** | 7.0 | 19.1 ± 0.4 | 15 ± 2 |
|  | 2.6 | 15.6 ± 0.7 | 11 ± 4 |
| **Extreme halophilic** | 7.0 | 12.7 ± 0.5 | 22 ± 2 |
|  | 2.6 | 12.9 ± 0.6 | 28 ± 4 |
| **Halotolerant** | 7.0 | 15.7 ± 0.4 | 17 ± 2 |
|  | 2.6 | 16.0 ± 0.5 | 20 ± 3 |

We also tested the effect of sulfate at lower pH (Table 2). For the non-halophilic and the intermediate-halophilic proteins, we find that the stabilizing effect is the same than that at neutral pH (Fig. 3A and 3C). In contrast, we observe that the sulfate-induced stabilizing effect for the most halophilic protein is higher at low pH than that at neutral pH (Fig. 3B). We speculate that this is because the neutralization of the carboxylic groups enables sulfate to explore the surface of the protein unconstrained from the electrostatic repulsion of carboxylates, enhancing its stabilizing effect. In other words, the negatively charged carboxylates “shield” the protein from the sulfate anion, and thus reduce the sensitivity of the protein to its stabilizing effect. We note, however, that the poor solubility of the extreme halophilic protein at these conditions (i.e., low pH and high concentrations of sulfate) reduces the precision of our measurements and complicates the interpretation of the results.

## Discussion

The study of extremophile adaptation has attracted a significant attention from the scientific community. Given the extreme physicochemical constraints that these organisms face, studying their adaptive mechanisms at a molecular level can help us understand the fundamental biophysical principles that regulate biomolecular function (Vallina Estrada and Oliveberg, 2022). For halophilic adaptation in particular, this interest arises in great part because it can help understanding the fundamental biophysical contributions to the interaction between biomolecules and the solvent, and how solvent composition affect the function, stability and structure of biomolecules (Ortega et al., 2020; Cuevas-Velazquez et al., 2021). In addition to this, halophilic adaptation is particularly attractive because, in contrast to other extremophile adaptations, it features a very characteristic fingerprint in the amino acid composition. As such, halophilic proteins are a convenient model system to understand the effect of solvent composition of protein stability, solubility and activity.

We present here a detailed characterization of the folding thermodynamics of model proteins with different degrees of halophilicity. In doing this, we find that halophilic amino acids reduce the mesophilic stability of proteins under low salt conditions, in great part due to the electrostatic repulsion between the highly abundant negatively charged carboxylates. This halophilic amino acid signature, however, drives haloadaptation by improving stability and solubility at high salt concentrations. This trade-off starts paying off at potassium chloride concentrations above 1M, precisely the limit beyond which extreme halophilic adaptation switches to the mechanism based on intracellular salt accumulation. For example, in the absence of salt and at neutral pH, the extreme halophilic variant is 8.5 kJ mol^-1^ less stable than wild type protein L, but achieves the same (or higher) stability at potassium chloride concentrations above 1.5 M. Moreover, for the three proteins and at all the conditions tested here, the mesophilic stability correlates fairly well with the degree of potassium chloride-induced stabilization (Fig. 4). This indicates that, by engineering the amino acid sequence, haloadaptation tunes the stability of halophilic proteins under hypersaline conditions to achieve values comparable to those of their non-adapted mesophilic counterparts under normal saline conditions, converging in an optimal stability to maintain a functional protein homeostasis.

**Figure 4.**
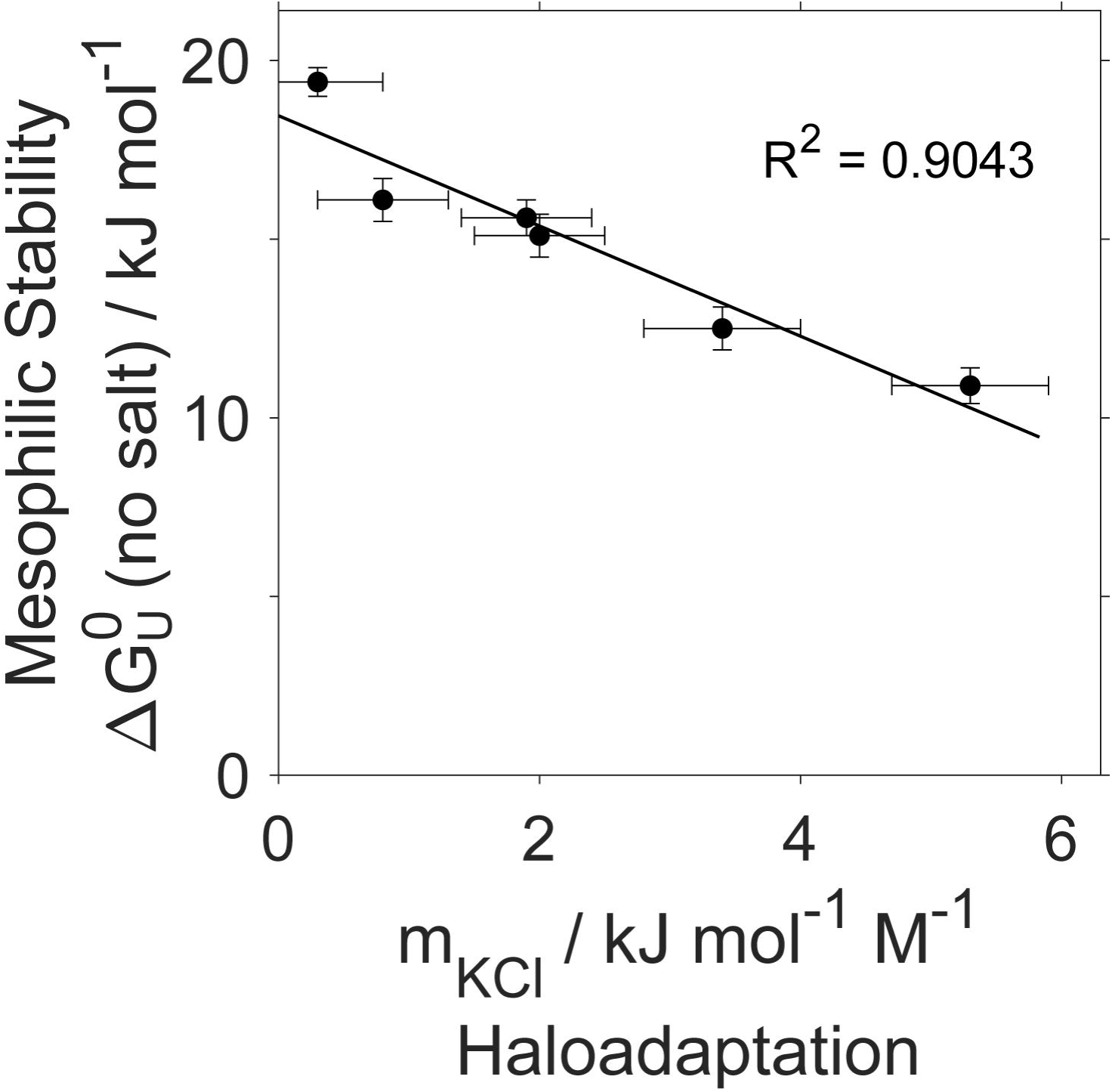
Trade-off between stability and salt-induced stabilization. For the three proteins and at all the conditions tested here, the mesophilic stability at low salt concentrations correlates with the stabilizing effect induced by potassium chloride. This suggests that halophilic adaptation engineers the surface of halophilic proteins to achieve an optimal thermodynamic stability at a given solvent composition.

We have also evaluated the role of electrostatics in haloadaptation. By measuring the effect of salt on the thermodynamics of folding at different pH values, we have accurately quantified the effect of the electrostatic interactions between the protein’s carboxylates and the ions in solution. Our results show that, while electrostatic interactions favor the stability of halophilic proteins in environments with high salt concentrations, they only account for a minor fraction of the total salt-induced stabilization. More specifically, for our extreme halophilic model these interactions account for 2 kJ mol^-1^ M^-1^, which is merely about one third of the total stabilizing effect of potassium chloride. Further supporting this, even when we neutralize all carboxylates at low pH we still observe a higher stabilizing effect for the extreme and intermediate halophilic variants compared to the non-halophilic protein, both with potassium chloride and with sodium sulfate. Our results thus suggest that halophilic adaptation also optimizes other aspects of amino acid composition besides electrostatic interactions, such as excluded volume and preferential cosolute exclusion (Schellman, 1990; Tadeo et al., 2009; Graziano, 2012; Graziano and Merlino, 2014; Ortega et al., 2015). To that end, substituting lysines for short and polar amino acids, such as glutamate, aspartate and glutamine, is particularly well-suited because it introduces negative charges while simultaneously reducing the apolar content of the surface. This is consistent with experimental studies showing that the interactions of ions with macromolecular surfaces are largely driven by weak, dispersive forces (Gurau et al., 2004; Chen et al., 2010); with computational studies suggesting that interactions between carboxylates, ions and water molecules in halophilic proteins are rather weak and unspecific, and similar to those of non-halophilic proteins (Geraili Daronkola and Vila Verde, 2021; Geraili Daronkola and Vila Verde, 2023); and with biological evidence showing that acidic proteomes can also adapt to low intracellular potassium chloride concentrations (Deole et al., 2013).

In our prior work, we studied the effect of different salts on the stability, structure, dynamics, and catalytic activity of various halophilic proteins (Tadeo et al., 2009; Tadeo et al., 2009; Ortega et al., 2011; Qvist et al., 2012; Ortega et al., 2015). In doing this, we found that the reduction of the solvent-exposed hydrophobic area was a key driver for halophilic adaptation, and more relevant than the content of negatively charged residues (Tadeo et al., 2009). Based on those results, we proposed a mechanistic model in which, while part of the halophilic adaptation indeed stems from additive weak carboxylate-cation interactions, its most significant fraction stems from preferential hydration and ion exclusion (Ortega et al., 2015). By increasing the fraction of polar area, halophilic amino acids facilitate hydration, which enhances ion exclusion from unfolded conformations. This imposes an energetic penalty associated with solvating separately ions and macromolecules, which becomes particularly costly at high salt concentrations where water is scarce. As a result, the unfolded state of halophilic proteins becomes much less favorable at high salt concentrations, thus producing a much higher stabilization of halophilic proteins. Our results showing that the higher salt-induced stabilization of halophilic proteins persists even in the absence of negatively charged carboxylates are consistent with these mechanistic hypotheses, and allow us to quantify accurately each contribution.

Our results also show the importance of electrostatic interactions for the solubility of halophilic proteins in high salt environments. We have observed that neutralization of carboxylates at low pH leads to a decrease in the solubility of both extreme and intermediate halophilic proteins. While at neutral pH the halophilic variants remained soluble above 1.25 M potassium chloride, at low pH we observed protein aggregation at potassium chloride concentrations beyond 0.75 M (Fig. 2). This is in contrast with the behavior of wild type protein L, which remained soluble at potassium chloride concentrations well above 1 M both at neutral and low pH. Similarly, in the presence of sodium sulfate we also observed a reduction in solubility at low pH compared to neutral pH conditions for the two halophilic proteins (Fig. 3). These qualitative observations are thus consistent with evolutionary adaptation favoring negatively charged amino acids to improve solubility at the highly crowded and hypersaline cytoplasm of halophilic organisms. To that end, glutamates and aspartates are particularly well suited, owing to their short, polar and negatively charged side chains.

In summary, our results provide a quantitative description of the extent to which interactions between negatively charged residues on the surface of halophilic proteins with potassium cations are important in halophilic adaptation. While these interactions seem to be significant in halophilic proteins, in light of our findings the magnitude of these electrostatic effects in halophilic adaptation has probably been overestimated in the past, in detriment of other contributions, such as excluded volume, hydrophobic effect, and preferential ion exclusion from the protein surface.

## Conclusions

We have studied the role of electrostatics in halophilic adaptation by measuring the effect of salts on the stability of proteins with different degrees of halophilicity. More specifically, we have studied a non-halophilic protein and two variants thereof engineered to feature intermediate and extreme halophilic amino acid sequences. We have then measured the effect of two different salts on the stability of these three proteins: potassium chloride, the most abundant ionic specie in the cytoplasm of extreme halophilic organisms, and sodium sulfate, one of the most stabilizing salts in the Hofmeister series. We have performed these measurements at a neutral pH, but also at a pH low enough to protonate all carboxylic groups present in the proteins. Our results show a thermodynamic trade-off by which the decrease in stability under mesophilic conditions is compensated by an enhanced stabilization at high salt concentrations. Haloadaptation thus optimizes the stability of halophilic proteins at hypersaline conditions to match that of mesophilic proteins under normal saline conditions. In addition to this, our experimental design has allowed us to separate the effect of carboxylate-potassium electrostatic interactions from the overall effect of potassium chloride on protein stability. By doing this, we find that carboxylate-related electrostatic interactions only account for about one third of the stabilizing effect of potassium chloride on halophilic proteins. Our results are consistent with a model in which the salt-induced stabilization of halophilic proteins, and thus haloadaptation, results mainly from preferential ion exclusion and competitive hydration, with electrostatic interactions being important for stability, albeit to a lesser extent, and for solubility.

## Experimental methods

### Protein expression and purification

We used the B1 domain of Protein L from *Finegoldia magna* (formerly *Peptostreptococcus magnus*) as our model protein. The amino acid sequence is MGSEEVTIKA NLIFANGSTQ TAEFKGTFEK ATSEAYAYAD TLKKDNGEWT VDVADKGYTL NIKFAG. In addition, we also used two variants in which we substituted the five lysines at positions 23, 28, 42, 54 and 61 by glutamates to engineer an extreme halophilic variant, and by glutamines for an intermediate halotolerant variant. We introduced the gene sequences into a pET16b plasmid (Novagen) between NcoI and XhoI restriction sites (GenScript), and then transformed BL21(DE3) Escherichia coli (New England BioLabs) with this plasmid. We grew the cell in LB media until an OD600 of 0.6, induced protein expression by adding 1 mM IPTG and continued incubation at 37°C for 4 hours. After harvesting the cells by centrifugation, we purified the proteins following previously described protocols (Tadeo et al., 2009). Briefly, we lysed the cells by sonication in lysis buffer containing 20 mM sodium phosphate at pH 6.0. We separated the lysate by centrifugation, subjected it to a heat shock treatment at 75°C for 5 minutes, allowed to recover room temperature under gentle shaking, and centrifuged to remove denatured proteins and cellular debris. We then further purified the supernatant by size exclusion chromatography on a HiLoad 26/600 Superdex 75 pg column (GE Healthcare) pre-equilibrated with the same buffer. When necessary to remove nucleic acid contamination, we also performed ion-exchange chromatography. To do this, we loaded the protein solution onto a HiTrap Q FF column (GE Healthcare) and eluted it with a linear gradient of 0–1 M NaCl in 20 mM sodium phosphate at pH 6.0. We assessed protein purity by SDS-PAGE and determined the protein concentration by measuring absorbance at 280 nm.

### Chemical denaturation experiments

We performed chemical denaturations using guanidinium chloride, which we monitored by circular dichroism (CD) on a J-850 spectrophotometer (JASCO, MD) and fluorescence spectroscopy. We prepared a 4 µM protein (0.4 µM for the extreme halophilic protein for the experiments at pH 2.6, due to its lower solubility) solution in 20 mM sodium phosphate buffer at pH 7.0, or 20 mM glycine-hydrochloride buffer at pH 2.6, and the desired concentration of potassium chloride or sodium sulfate. For the sodium sulfate experiments, we included a background of 0.25 M potassium chloride to compensate non-linear effects of guanidinium chloride at low ionic strengths. We then titrated it with a guanidinium chloride solution in a quartz cuvette of 1 cm path length at 25°C. After each guanidinium chloride addition, we stirred the sample for 1 minute before measurement to allow for sufficient equilibration. We recorded circular dichroism spectra from 215 to 250 nm at 50 nm/min with a bandwidth of 4 nm, digital integration time of 4 s, and accumulating 3 scans. We used changes in ellipticity at 222-230 nm to monitor the protein unfolding equilibrium.

### Data analysis

We analyzed the data using custom Matlab scripts to fit the denaturation curves to a two-state folding equilibrium model following the linear extrapolation method (Santoro and Bolen, 1988). The circular dichroism ellipticities were fit to the following equations:

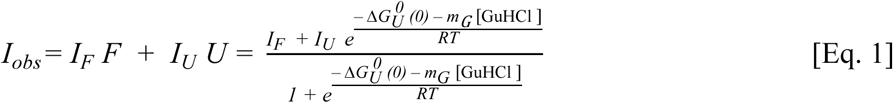

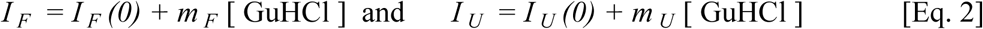

We used a non-linear least-squares minimization method to fit the data and extract six parameters: the intrinsic ellipticities for the folded and unfolded states in the absence of denaturant, *I_F_(0)* and *I_U_(0)*, the slopes for the linear dependence of the ellipticities on guanidinium chloride concentration, *m_F_* and *m_U_*, and the thermodynamic parameters for the unfolding free energy in the absence of denaturant, *11G^0^_U_(0)*, and the slope of the linear dependence of the unfolding free energy with the denaturant concentration, *m_G_*. Uncertainties reported correspond to 95% confidence intervals derived from the fit.

### Data fitting

To analyze the effect of potassium chloride and sodium sulfate on protein stability, we fitted the data to the following linear equation:

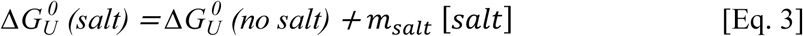

We performed a linear regression treating cosolute concentration as the independent variable and protein stability as the dependent variable using the Matlab “*polyfit*” function. The slope of this linear fit, *m_salt_*, represents the effect of the cosolute on protein stability, with a positive slope indicating a stabilizing effect (increasing stability with increasing cosolute concentration), while a negative slope indicates a destabilizing effect (decreasing stability with increasing cosolute concentration). In turn, the intersect, 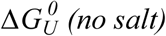, represents the mesophilic stability, which is the intrinsic stability of the protein at low salt concentrations. To calculate the associated errors, we consider the individual uncertainties for each stability measurement. To do this, we use a Monte Carlo approach to generate 1,000 new datasets in which we propagate the errors for each data point assuming a normal distribution, and fit each one of these to a linear equation. The errors reported correspond to 95% confidence intervals of the distributions of the slopes and intersects derived from these fits.

